# MicroRNAs provide the first evidence of genetic link between diapause and aging in vertebrates

**DOI:** 10.1101/017848

**Authors:** Luca Dolfi, Mario Baumgart, Marco Groth, Roberto Ripa, Matthias Platzer, Alessandro Cellerino

## Abstract

Diapause and aging are controlled by overlapping genetic mechanisms in *C.elegans* and these include microRNAs (miRNAs). Here, we investigated miRNA regulation in embryos of annual killifish that naturally undergo diapause to overcome desiccation of their habitats. We compared miRNA expression in diapausing and non-diapausing embryos in three independent lineages of killifish. We identified 13 miRNAs with similar regulation in all three lineages. One of these is miR-430, which is known as key regulator of early embryonic development in fish. We further tested whether this regulation overlaps with the aging-dependent regulation of miRNAs in one annual species: *Nothobranchius furzeri*. We found that miR-101a and miR-18a are regulated in the same direction during diapause and aging. These results provide the first evidence that overlapping genetic networks control diapause and aging in vertebrates and suggest that diapause mimics aging to some extent.

---

*Diapause* is a suspension of larval development to overcome adverse conditions. An overlap in the genetic mechanisms controlling diapause (dauer) and longevity was discovered in *C. elegans* and led to the identification of insulin/IGF pathways as a conserved regulator of aging (Kenyon 2011). MicroRNAs (miRNAs) are part of the genetic network that regulates longevity and diapause (de Lencastre *et al.* 2010; Pincus *et al.* 2011; Zhang *et al.* 2011). These results prompted us to investigate a possible overlap in microRNA regulation during aging and diapause in vertebrates.

Diapause is a survival strategy in *annual* fish that are adapted the alternation of wet- and dry-season in Africa and South America (Wourms 1972) and is present in three independent evolutionary lineages of killifishes as the result of multiple events of loss and re-gain of an ancestral annual life history (Murphy & Collier 1997; Hrbek & Larson 1999). In each of these clades, annual species have as nearest neighbor a non-annual species (Fig. 1A) and this *taxon* offers a unique paradigm to test parallel evolution of life-history traits.

**Figure 1 (A).**
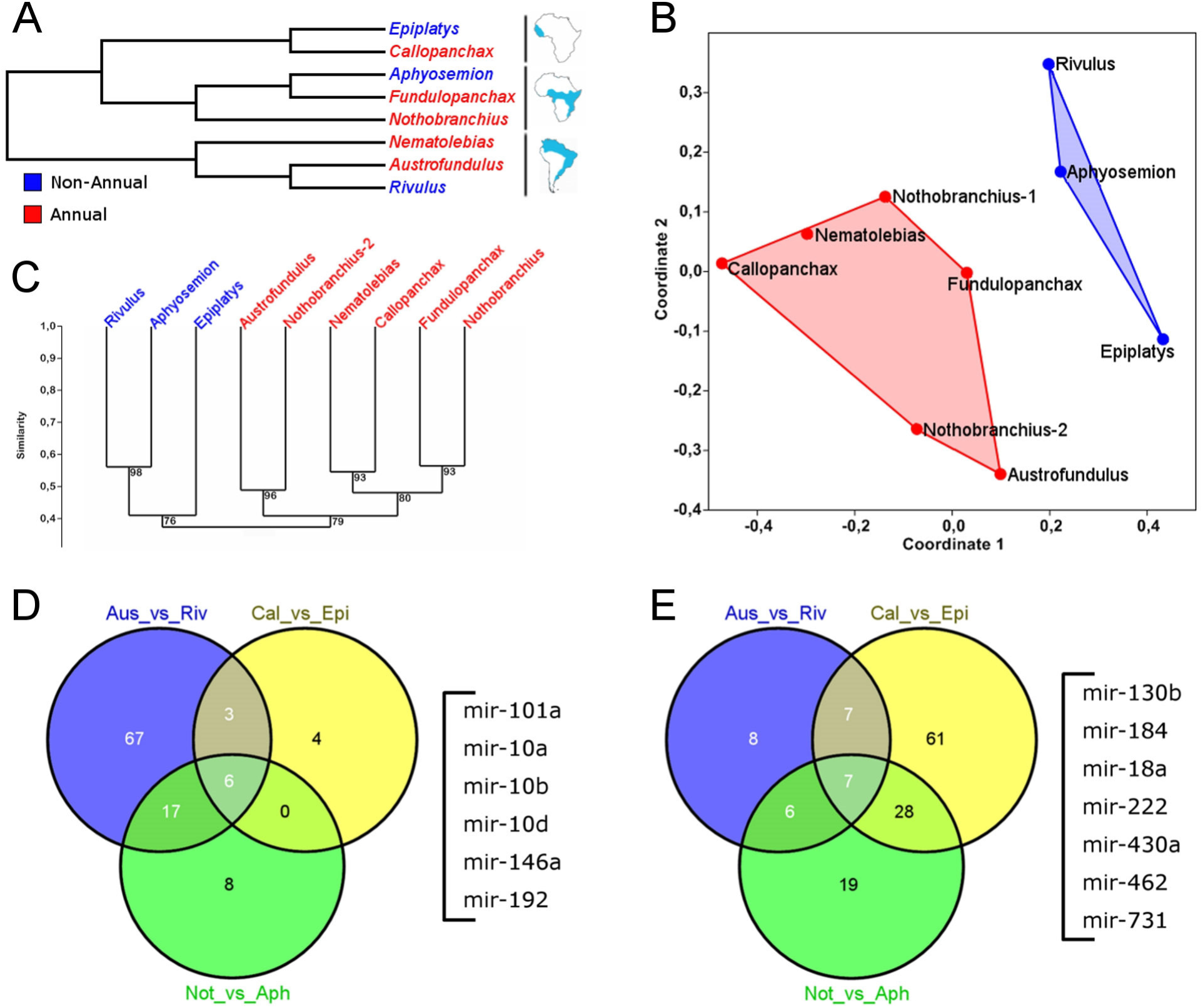
Cladogram of killifish depicting the species used in this study. Red branches represent annual *genera* and blue branches non-annual *genera.* (B) MDS plot of all analyzed samples. The red dots represent annual *genera* and blue dots non-annual *genera. Nothobranchius*-1 and -2 are biological replicates. (C) Hierarchical clustering of the same samples as in (B), Bray-Curtis dissimilarity index. (D,E) Venn plot of common up-and down-regulated miRNAs, respectively. Each contrast was performed between annual- and non-annual species of the same evolutionary lineage. Species codes: Nfu = Nothobranchius furzeri, Ast = Aphyosemion striatum, Coc = Callopanchax occidentalis, Eda = Epiplatys dageti, Ale = Austrofundulus leohoignei, Rcy = Rivulus cylindraceus.

Aim of the present study was to characterize the miRNA signature of diapause in annual fish and compare it with the signature we previously detected during aging of the annual species *Nothobranchius furzeri* (Baumgart et al. 2012).

We used miRNA-Seq (Morin *et al.* 2008) to identify and quantify expression of miRNAs during diapause. We analyzed diapausing embryos and embryos in the corresponding developmental stage (mid-somite) for five annual species and three non-annual species from all three lineages (Fig.1A). We used hierarchial clustering (Fig. 1C) and multi dimensional scaling (MDS, Fig.1B) to visualize global aspects of miRNA regulation. Results showed that samples cluster according to their physiological status (diapausing or not) and not to their phylogenetic relationships. Similarly, diapausing and non-diapausing samples occupy non-overlapping regions of the MDS space.

We identified miRNAs differentially expressed between closely related annual and non-annual species common to all three lineages (Kal’s Z-test, FDR-corrected p<0.01 and absolute fold change > 1.5). Venn analysis revealed six miRNAs that were up-regulated in diapausing embryos in all three lineages (Fig 1D, Table S1): miR-10a/b/d, miR-101a, miR-146a and miR-192. Seven miRNAs were down-regulated in diapausing embryos in all three lineages (Fig 1E, Table 1): miR-130, miR.184, miR-18a, miR-22a, miR-430a, miR-462, and miR-731. MiR-430 controls transition from maternal to zygotic transcription in fish eggs (Giraldez *et al.* 2006). This transition appears normal however in annual fish and well before diapause I, as assessed by onset of paternal transgene expression in *N. furzeri* (Fig. 2), suggesting a novel role for miR-430.

**Table 1.** Fold-changes of miRNAs that were detected as differentially-expressed in all three lineages. The miRNA name indicates the *D. rerio* miRNA that was used for annotation. Species codes: Nfu = *Nothobranchius furzeri*, Ast = *Aphyosemion striatum*, Coc = *Callopanchax occidentalis*, Eda = *Epiplatys dageti*, Ale = *Austrofundulus leohoignei*, Rcy = *Rivulus cylindraceus*.

| miRNA | Fold-change |  |  |  |
| --- | --- | --- | --- | --- |
|  | Nfu vs. Ast | Coc vs. Eda | Ale vs. Rcy | Average |
| up-regulated |  |  |  |  |
| miR-10a | 5,76 | 4,53 | 8,91 | 6,4 |
| miR-10b | 3,39 | 2,05 | 6,78 | 4,07 |
| miR-10d | 2,66 | 1,99 | 19,39 | 8,01 |
| miR-101a | 4,82 | 2,39 | 18,93 | 8,71 |
| miR-146 | 1,93 | 6 | 5,08 | 4,34 |
| miR-192 | 7,7 | 6,92 | 23,26 | 12,63 |
| down-regulated |  |  |  |  |
| miR-130b | -1,58 | -2,39 | -91,7 | -31,89 |
| miR-184 | -2,81 | -7,83 | -1,52 | -4,05 |
| miR-18a | -2,07 | -14,96 | -1,64 | -6,22 |
| miR-222a | -3,6 | -14,67 | -2,17 | -6,81 |
| miR-430a | -3,72 | -5,24 | -35,53 | -14,83 |
| miR-462 | -17,27 | -22,53 | -1,57 | -13,79 |
| miR-731 | -17,06 | -47,39 | -4,24 | -22,9 |

**Figure 2.**
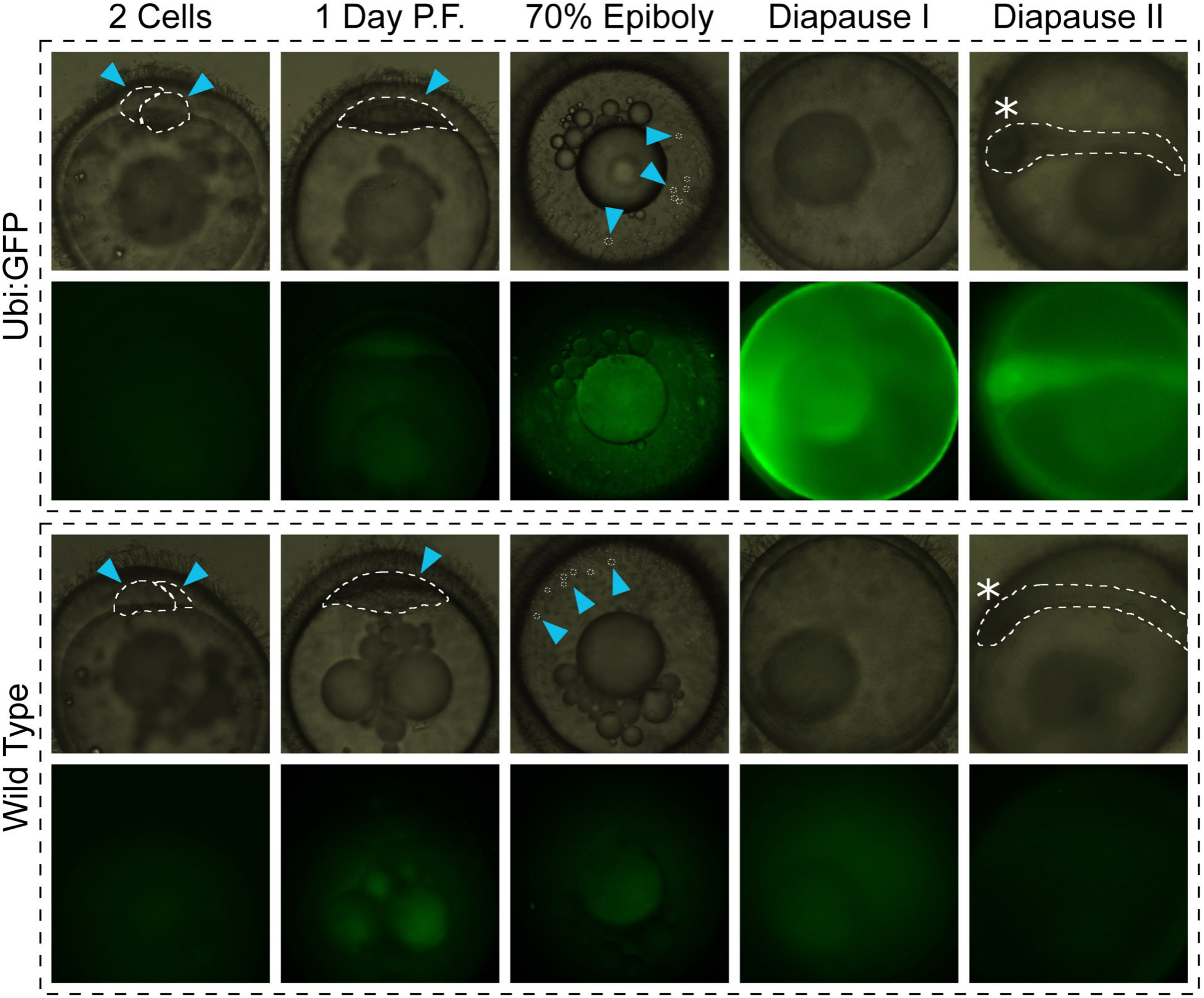
Expression of paternal transgene in *N. furzeri*. The expression of paternal genes is shown in a transgenic line expressing eGFP under the ubiquitin promoter. The top two rows illustrate a transgenic egg and the bottom two row illustrate a control from the same clutch. The dotted lines indicate the boundary of the embryo. Dispersed cells are shown in the dispersed state (70% epiboly). Please note that the first expression of the transgene is observed at the dome stage when mid-blastula transition is expected.

In *C. elegans,* some miRNAs are regulated in similar directions during diapause and aging. In particular miR-71, that is induced during diapause and is necessary for survival of diapausing *larvae* (Zhang et al. 2011). Expression of miR-71 increases with age and drops late in life (Pincus *et al.* 2011). Deletion of miR-71 results in shortened life span (de Lencastre *et al.* 2010) and the timing of miR-71 drop is predictive of individual worm lifespan (Pincus *et al.* 2011). In the annual species *N. furzeri*, we detected four miRNAs that are up-regulated during aging in brain liver and skin (Baumgart *et al.* 2012). One of these miRNAs, miR-101a, is also up-regulated during diapause in all three annual lineages analyzed here. In human cancer cells, miR-101 represents a key node of a regulatory network with transcription factors and epigenetic modulators as the first neighbors and genes involved in cell-cycle progression as second neighbors (Huang *et al.* 2012). In addition, we detected conserved down-regulation of miR-18a during diapause. This miRNA is a member of the miR-17^∼^92 genomic cluster, also known as oncomiR-1, that plays a major role in promoting cell proliferation in cancer cells (Olive *et al.* 2010). These miRNAs are down-regulated during aging of human mitotically active cells (Grillari *et al.* 2010) and in brain and skin of *N. furzeri* (Baumgart *et al.* 2012). Regulation of miR-18a and miR-101a are therefore most likely related to mitotic arrest observed in diapausing killifish embryos (Meller *et al.* 2012). It is an interesting observation that the regulation of these miRNAs during diapause follows the same direction observed during aging and mirrors the regulation of miR-71 in *C. elegans*.

These results provide the first evidence that overlapping genetic networks control diapause and aging in vertebrates.

## Experimental Procedures

### Fish raising

All the fishes used in this work were raised in 35 lt tanks (25cm × 50cm × 28cm) filtered with an air-driven sponge filter at these densities:

- *Aphyosemion australe, Aphyosemion striatum, Nothobranchius furzeri*, *Nothobranchius guentheri*, *Nothobranchius korthause, Scriptaphysemion guignardi* and *Epiplatys dageti:* 3 or 4 couples each tank;
- *Nothobranchius melanospilus:* 1 or 2 couples each tank;
- *Callopanchax occidentalis:* 1 couple each tank;
- *Rivulus cylindraceus, Rachovia brevis:* 3 couples each tank

Water parameters used for all species

- pH: 7-8;
- KH: 3-5;
- T: 23-25 °C

25% of the water in each tank was replaced once a week.

### Fish feeding

All the fishes were fed with granular food (Microgranuli, SHG^®^, Alessandria, Italy) twice a day and newly-hatched brine-shrimps (Premium Artemia, SHG^®^, Alessandria Italy) once a day (rising to three times a day will result in increased eggs production) for the amount they can eat in ten minutes.

### Fish breeding

#### Annual and non-annual fishes were bred using two different methods

For annual fishes, a bowl (from 20cm × 15cm × 10cm to 9cm × 9cm × 4cm), half-full of river sand (**ø** 0.3mm), was put on the bottom of the tank. More than one small bowl (0.7-1.3 for each male in the tank) worked better in tanks with more than 4 males. After a short period, or after a short conditioning (see below), fishes learned to go in the bowl and breed on the surface of the sand, laying and burying eggs deep into it.

Eggs could then be collected by removing the bowl from the tank, sieving the sand with a sieve (1mm grid) and then trasferring the eggs from the sieve to a Petri dish previously filled with tap water. Non-annual fishes were bred differently. 25cm long “breeding mops” were made using 2mm thick green/brown/grey wool (100% acrylic), using a Bijou (by Sigma-Aldrich^®^) screwed into the top of each mop to make them floating. One or more mops (optimum is one each two males) were put in the tanks. After a short period, or after a short conditioning (see below), fishes learned to go in the mops and breed inside it, laying eggs on the wool filaments. Eggs could then be collected by removing the mops from the tanks and inspecting them, eggs were removed with one finger tips (eggs are adhesive and hard) and putting them into a Petri dish previously filled with tap water.

Conditioning for breeding

1. To force the breeding time is often useful a relative short period of habituation (from 5 days to 2 weeks), during which fish are put in a comfortable situation for coupling (with sand on the bottom of the tank or mops on the top), for a short period (1.5 to 3 hours) daily, at the same time of the day.
2. Intense breeding sessions (breeding fish every day for more than 5 days) would result in decreasing the amount of eggs produced. This can be solved by 1 or 2 days of rest and it is recommended to apply the period of rest before any significant experiments. Notice that if the non-breeding period is prolonged too much it can result in impaired eggs production when breedings are restored.
3. Fishes can be forced to produce eggs by separating males from females for several days (from 5 days to 2 weeks) and then putting them together in a tank with plenty of mops/sands.
4. The best setup for breeding is a 5-25 lt (depending on species) tank with one male, two females and one mop/sand. This method works better in combination with point 3 while is often in conflict with point 1.
5. *Rivulus cylindraceus* breeding: *Rivulus cylindraceus* were able to produce eggs only after at least 10 days of gender separation, during which 4 males and 4 females are put into 2 different 40+ lt tanks set up with bricks where they can hide. Afterwards, they were put all together again in a 3 lt tank with 5 mops.

### RNA Isolation

For each species 20 eggs were pooled for each preparation. Total RNA was isolated using QIAzol (Qiagen) according to the manufacturer’s protocol. 1 ml cooled QIAzol and one 5 mm stainless steel bead (Qiagen) was added. Homogenization was performed using a TissueLyzer II (Qiagen) at 20 Hz for 2-3×1 min. After incubation for 5 min at room temperature 200 μl chloroform was added. The tube was shaken vigorously for (at least) 15 s and incubated for 3 min at room temperature. Phase separation was achieved by centrifugation at 12,000×g for 20 min at 4°C. The aqueous phase was transferred into a fresh cup and 10 μg of Glycogen (Invitrogen, Darmstadt, Germany), 0.16×volume NaAc (2 M; pH 4.0) and 1.1×volume isopropanol were added, mixed thoroughly and incubated for 10 min at room temperature. The RNA was precipitated by a centrifugation step with 12,000 × g at 4°C for 20 min. The supernatant was removed and the pellet was washed with 80% Ethanol twice and air dried for 10 min. The RNA was resuspendet in 20 μl DEPC-treated water by pipetting up and down, followed by incubation at 65°C for 5 min. The RNA was quantified with a NanoDrop 1000 (PeqLab, Erlangen, Germany) and stored at -80°C until use.

### Small RNA Sequencing

For small RNA sequencing, small RNA cDNA libraries were prepared as follows: for each developmental stage equal quantities (1 µg) of total RNA were submitted independently to Illuminas TruSeq small RNA sample preparation protocol (Illumina Inc., San Diego, USA). In brief, the library preparation was performed as follows: RNA was ligated with proprietary adapters to the 5’ and 3’ termini of the RNA. The adapter ligated samples were used as templates for cDNA synthesis. The cDNA was amplified with 13 PCR cycles to produce sequencing libraries, introducing specific nucleotide codes for indexing of libraries, to allow multiplexed sequencing. cDNAs were purified by 10% Novex TBE polyacrylamide gel electrophoresis (Invitrogen) and eluted into 300 μl elution buffer (Illumina) for at least 2 hours at room temperature, to enrich for molecules containing inserts in the range of 18–33 nt. The resulting gel slurry was passed through a Spin-X filter (IST Engeneering Inc., Milpitas, CA, USA) and precipitated by the addition of 20 μg glycogen, 30 μl of 3M NaOAc, pH5.2, and 975 μl of pre-chilled (-20°C) ethanol. After washing with 70% ethanol, the pellet was dryed at 37°C for 5-10 min and dissolved in 10 μl resuspension buffer (Illumina). The purified libraries were quantified on the Agilent DNA 1000 chip, diluted to 10 nM and subjected to sequencing-by-synthesis on Illumina HiSeq 2000.

## Analysis of sequencing data

Individual sequence reads with base quality scores were produced by Illumina sequencing. The data was analyzed by the use of CLC-worbench 4 (CLCbio, Arhus, Denmark). After eliminating reads with low quality and trimming the 3’ adaptor sequence, the remaining 18- to 33-nt reads represented by at least 30 sequence reads were grouped into unique sequence clusters. Annotation of sequence clusters was performed, allowing 2 mismatches and up to 2 additional bases at each end, by using 4 fish species as references from miRBase v16.0: *Danio rerio*, *Tetraodon nigroviridis*, *Fugu rubripes*, *Oryzias latipes*. Statistical significance was assessed using Kal’S Z test as implemented in CLC-workbech setting a p-adjusted value of 0.05.

Hierarchical clustering and multi-dimensional scaling were performed using PAST (Hammer *et al.*, 2001). Venn diagrams were generated using Venny (Oliveiros, 2007)

## Acknowledgments

This work was partially supported by an internal grant of Scuola Normale Superiore and by the German Ministry for Education and Research (JenAge; BMBF, support code: 0315581A, 0315581C)

